# Deep Generative Models for 3D Compound Design

**DOI:** 10.1101/830497

**Authors:** Fergus Imrie, Anthony R. Bradley, Mihaela van der Schaar, Charlotte M. Deane

## Abstract

Rational compound design remains a challenging problem for both computational methods and medicinal chemists. Computational generative methods have begun to show promising results for the design problem. However, they have not yet used the power of 3D structural information. We have developed a novel graph-based deep generative model that combines state-of-the-art machine learning techniques with structural knowledge. Our method (“DeLinker”) takes two fragments or partial structures and designs a molecule incorporating both. The generation process is protein context dependent, utilising the relative distance and orientation between the partial structures. This 3D information is vital to successful compound design, and we demonstrate its impact on the generation process and the limitations of omitting such information. In a large scale evaluation, DeLinker designed 60% more molecules with high 3D similarity to the original molecule than a database baseline. When considering the more relevant problem of longer linkers with at least five atoms, the outperformance increased to 200%. We demonstrate the effectiveness and applicability of this approach on a diverse range of design problems: fragment linking, scaffold hopping, and proteolysis targeting chimera (PROTAC) design. As far as we are aware, this is the first molecular generative model to incorporate 3D structural information directly in the design process. Code is available at https://github.com/oxpig/DeLinker.

## Introduction

Drug design is an iterative process that requires potential compounds to be optimised for specific properties, ranging from binding affinity to pharmacokinetics. This process is challenging, in part due to the size of the search space^1^ and discontinuous nature of the optimisation landscape.^2^ Typically molecule design is undertaken by human experts, and therefore is a subjective process.

Machine learning models for molecule generation have been proposed as an alternative to human-led design and rules-based transformations.^3–5^ Generative models have adopted either the SMILES string representation of molecules^6–10^ or, more recently, graph representations.^11–15^ Existing generative models have primarily been used in two ways. First, methods have been developed to generate molecules that follow the same distribution as the training set, whether a general set of molecules^10^ such as ZINC^16^ or ChEMBL,^17^ or a more fo-cussed one such as inhibitors for a particular protein target.^7,18^ Second, generative models have been proposed to perform molecular optimisation, taking an input molecule and attempting to modify one, or several, chemical properties, typically subject to a similarity constraint.^15^

While substantial progress has been made for these two problems, current methods have inherent limitations, in particular for structure-based design. Only one approach to date has attempted to include any three dimensional (3D) information in the generative process,^19^ despite its importance for designing potent and selective compounds. In this work, 3D information was only provided implicitly to the generative model, and the method did not allow further control over generated compounds. ^19^ As a result, the generative model frequently changes the entire molecule. This is undesirable in many practical settings, such as the design problems described below.

Fragment-based drug discovery (FBDD) has become an increasingly important tool for finding hit compounds, in particular for challenging targets and novel protein families. FBDD utilises smaller than drug-like compounds (typically *<*300 Daltons) to identify low potency, high quality leads, that are then matured into more potent, drug-like compounds. One common way of maturing fragments hits is through a linking strategy, joining fragments together that bind to distinct sites via a linker. It is crucial for successful fragment linking that a linker does not disturb the original binding poses of each fragment.^20,21^ Thus compound suggestions have strong 3D constraints, determined by the binding mode of the fragments.

Scaffold hopping, though a distinct problem, shares some characteristics with fragment linking. The aim of scaffold hopping is to discover structurally novel compounds starting from a known active compound by modifying the central core structure of the molecule.^22^ Such a change can result in much improved molecular properties, such as solubility, toxicity, synthetic accessibility, affinity, and selectivity. ^22,23^

Numerous computational methods have been proposed for fragment linking or scaffold hopping.^24–29^ However, almost all methods published to date rely exclusively on a database of candidate fragments from which to select a linker, with the differences between approaches arising solely from how the database is searched, how the linked compounds are scored, or the contents of database itself. As a result, these methods are inherently constrained to a set of predetermined rules or examples, limiting exploration of chemical space. In addition, they can only incorporate additional structural knowledge (e.g. the fragment’s binding mode) via filtering or search mechanisms.

Current machine learning-based molecule generation methods are not suitable for the design tasks of fragment linking and scaffold hopping. These scenarios require proposed molecules to contain specific substructures, with the goal to design a molecule that maintains the binding mode of the original compound or fragments. Neither of these requirements have been explicitly included in previous methods.

In this work, we introduce the first graphbased deep generative method that incorporates 3D structural information directly into the design process. Our method takes as input two molecular fragments and designs a molecule incorporating both substructures, either generating or replacing the linker between them. This allows our method to handle structure-based design tasks such as fragment linking and scaffold hopping effectively. The generation process is protein context dependent, and integrates 3D structural information, specifically the distance between the fragments and their relative orientations. This 3D information is vital to successful compound design, and we demonstrate the limitations of omitting such information, both quantifying its impact in large-scale assessments and empirically showing how our model uses the structural information.

We first demonstrate the effectiveness of our proposed deep generative approach over a database method through large-scale computational assessments. We show that our method, DeLinker, designs 60% more compounds with high 3D similarity to the original molecule compared to a database-based approach on an independent test set. DeLinker outperforms the database approach by 200% when the evaluation is restricted to linkers with at least five atoms. We then apply our method to several case studies encompassing fragment linking, scaffold hopping, and PROTAC design. DeLinker frequently recovers the experimental end-point, even in cases where the linker was not present in the training set, and produces many novel designs with high 3D similarity to the original molecules.

## Methods

The method takes two fragments and their relative position and orientation and generates or replaces the linker between them. This is achieved by building new molecules in an iterative manner “bond-by-bond” from a pool of atoms that can be initialised with partial structures (Figure 1). In this framework, the user is able to control the generation process by specifying both the substructures that should be linked and the maximum length of linker between them. In addition, 3D structural information in the form of the distance and angle between the starting substructures is provided to the model to inform the design process. Molecules are encoded by a set of 14 permitted atom types, and our model enforces simple atomic valency rules via a masking procedure to ensure chemical validity. This is the only chemical knowledge incorporated directly into our model; all other decisions required to generate molecules are learnt through a supervised training procedure.

**Figure 1:**
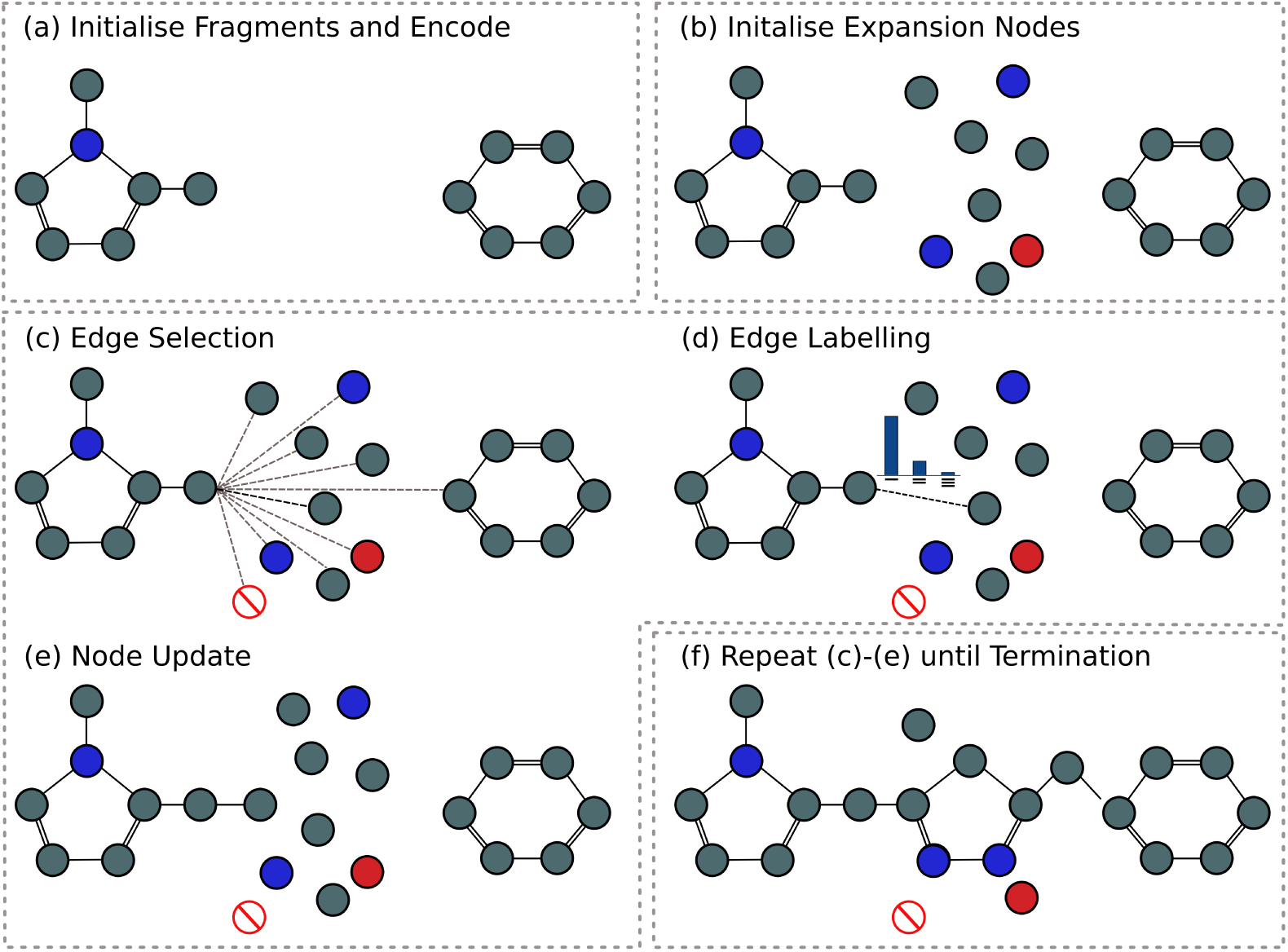
Overview of the generation process. The initial fragments (a) are iteratively expanded “bond-by-bond” (c)-(e) to produce a molecule including both fragments (f). Atoms are represented by nodes in a graph, with the colour of the nodes representing different atoms types, while bonds are represented by edges, with different edge types for single, double and triple bonds.

### Generative process

The generative process is illustrated in Figure 1 and is similar to Liu et al. ^14^ in that our method builds molecules “bond-by-bond” in a breadth-first manner. Generation is initialised with two fragments or substructures which are to be linked together with structural information providing the distance and angle between the substructures. The fragments are converted to a graph representation, where atoms and bonds are represented by nodes and edges, respectively. Each node is associated with a hidden state, *z*_*ν*_, and label, ***l***_*v*_, representing the atom type of the node. A list of the 14 permitted atom types can be found in the Supporting Information. The graph is passed through an encoder network, a standard gated graph neural network (GGNN),^30^ and the hidden states of the nodes are updated to incorporate their local environment (Figure 1a).

Next, a set of expansion nodes are initialised at random, with hidden states ***z***_*v*_ drawn from the *h*-dimensional standard normal distribution, 𝒩 (**0**, ***I***), where *h* is the length of the hidden state (Figure 1b). The nodes are then labelled with an atom type according to their hidden state, ***z***_*v*_, by sampling from the softmax output of a learned mapping *f* (***z***_*v*_). Here, *f* is implemented as a linear classifier but could be any function mapping a node’s hidden state to an atom type. The number of expansion nodes determines the maximum length of the linker, and is a parameter chosen by the user.

The new molecule is constructed from this set of nodes via an iterative process consisting of edge selection, edge labelling, and node update (Figure 1c-e). At each step, we consider whether to add an edge between one of the nodes, *v*, and another node in the graph. *v* is chosen according to a deterministic first-in-first-out queue that is initialised with the exit vectors of each fragment. When a node is connected to the graph for the first time, it is added to the queue. New edges are added to node *v* until an edge to the stop node is selected. The node then becomes “closed” with no additional edges with that node permitted.

All possible edges between the node *v* and other nodes in the graph are considered (Figure 1c), subject to basic valency constraints. A single-layer neural network assesses the candidate edges using a feature vector. The feature vector for the edge between node *v* and candidate node *u* is given by

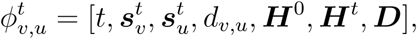

where 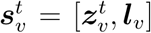 is the concatenation of the hidden state of node *v* after *t* steps and its atomic label, *d*_*v,u*_ is the graph distance between *v* and *u*, ***H***^0^ is the average initial representation of all nodes, ***H***^*t*^ is the average representation of nodes at generation step *t*, and ***D*** represents the 3D structural information. The feature vector provides the model with both local information for the node *v* and the candidate node *u* 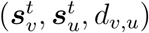, and global information regarding both the original graph specification (***H***^0^) and the current graph state (***H***^*t*^). The model is also provided with structural information (***D***), namely the relative distance and orientation between the starting substructures.

Once a node *u* has been selected, the edge between *v* and *u* is labelled as either a single, double, or triple bond (subject to valency constraints) by another single layer neural network taking as input the same feature vector 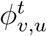 (Figure 1d).

Finally, the hidden states of all nodes are updated according to a GGNN (Figure 1e). At each step, we discard the current hidden states 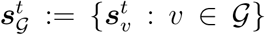 and and compute new representations 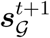 taking their (possibly changed) neighborhood into account. Note that 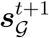 is computed from 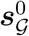 rather than 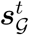. This means that the state of each node is independent of the generation history of the graph and depends only on the current state of the graph.

Steps c-e in Figure 1 are repeated for each node in the queue, until the queue is empty, at which point the generation process terminates. At termination (Figure 1f), all unconnected nodes are removed and the largest connected component is returned as the generated molecule.

### Multimodal Encoder-Decoder Setup

Our goal is to learn a multimodal mapping from unconnected fragments to connected molecules. During training, we utilised a data set of paired fragments and molecules and trained our model in a supervised manner to reconstruct known linkers. While in this data set there may be a unique molecule associated with two fragments, in practice there are many ways to link two fragments. As such, given a new pair of starting points, a model should be able to generate a diverse set of output compounds.

To this end, we took inspiration from Jin et al. ^15^ and augmented the basic encoder-decoder model with a low-dimensional latent vector ***z*** to explicitly encode the multimodal aspect of the output distribution. The generative mapping is converted from *F* : *X* ↦ *Y* to *F* : (*X,* ***z***) ↦ *Y*, where *X* represents the starting substructures and *Y* the connected molecule, with latent code ***z*** drawn from a prior distribution, chosen to be the standard normal distribution, 𝒩 (**0**, ***I***).

There are two challenges in learning this mapping. First, as shown in the image domain, ^31^ the latent codes are often ignored by the model unless they are forced to encode meaningful variations. Second, the latent codes should be suitably regularised so that the model does not produce invalid outputs. That is, the generated molecule *F* (*X,* ***z***) should belong to the domain of the target molecule *Y* (i.e. connected and able to satisfy the structural constraints provided) given a latent code drawn from the prior distribution. We overcome both of these challenges through our training procedure, where we derived ***z*** during training from the embedding of the linked molecule, but regularised the latent vector to follow a standard normal distribution so that we can sample ***z*** during generation.

### Training

We trained our generative model under a variational autoencoder (VAE) framework on a collection of fragment-molecule pairs (Figure 2). For a given pair of fragments *X* and linked molecule *Y*, the model is trained to reconstruct *Y* from (*X,* ***z***), while enforcing the standard regularisation constraint on both ***z*** and the encoding of *X*, ***z***_*X*_ := {***z***_*v*_ : *v* ∈ *X*}.

**Figure 2:**
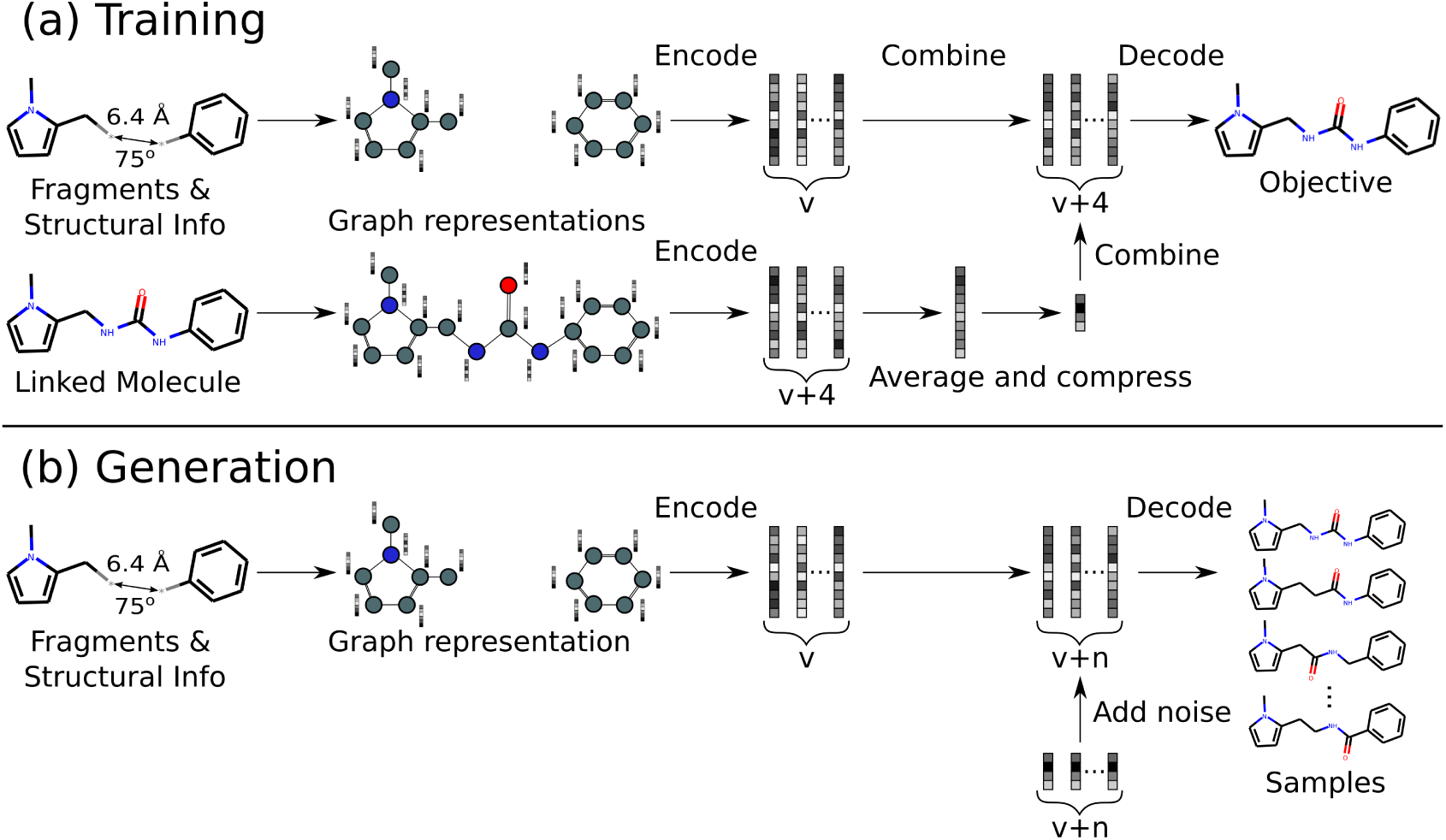
Illustration of training and generation procedures. (a) Pairs of fragments and linked molecules are provided as input. The model is trained to reproduce the linked molecule from a combination of the encodings of the fragments and linked molecule. (b) At generation time, the model is given only the unlinked fragments and structural information, and is able to sample a diverse range of linked molecules by combining the encoding of the fragments with random noise.

To encode meaningful variations, the latent code ***z*** is derived via a learnt mapping from the average of the node embeddings of the ground truth molecule *Y*, the linked molecule. Crucially, ***z*** is constrained to be a low dimensional vector to prevent the model from ignoring input *X* and degenerating to an autoencoder for *Y*. The decoder is trained to reconstruct *Y* when taking as input a combination of the low dimensional vector ***z*** and the node embeddings ***z***_*X*_ of the unlinked fragments *X* (Figure 2).

The training objective is similar to the standard VAE loss, including a reconstructionn loss and a Kullback-Leibler (KL) regularisation term:

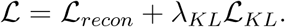

The reconstruction loss is composed of two cross-entropy loss terms, resulting from the error in predicting the atom types and in reconstructing the sequence of steps required to produce the target molecule.

The KL regularisation loss contains two terms, one for the encoding of the unlinked fragments *X*, the other for the low dimensional vector ***z*** derived from the linked molecule *Y*. These terms are the standard VAE terms capturing the KL divergence between the encoder distributions and the standard Gaussian prior.

We performed limited hyperparameter tuning, measuring performance via the validation loss and not generative performance directly. We found that overall the model was fairly robust to the choice of hyperparameters. Full details of the model architecture and hyperparameters can be found in the Supporting Information.

### Database method

Several traditional methods exist for linking fragments or replacing the core of a molecule.^24–29^ Almost all methods rely on a database from which to select linkers. As a baseline with which to compare our method, we created a set of all linkers from the training data and sampled from this set, joining the linker in one of the two possible orientations at random. This setup ensures that both methods are constructed using the same data, and allows us to assess whether the generated molecules have better shape complementarity than using linkers from the database, while still obeying 2D chemical constraints.

### Data sets

There have only been a limited number of examples of successful fragment linking or scaffold hopping reported. As such, for training and large scale evaluation, we constructed sets of fragment-molecule pairs using standard transformations from matched-molecular pair analysis.^5^

#### ZINC

To construct our training set, we used the subset of ZINC^16^ selected at random by Gómez-Bombarelli et al. ^10^ that contains 250 000 molecules. We constructed possible fragmentations of each molecule by enumerating all double cuts of non-functional group, acyclic single bonds, the same procedure adopted by Hussain and Rea ^5^. Fragmentations satisfying basic criteria regarding the number of atoms in the linker and fragments were retained, removing trivial and unrealistics scenarios (see Supporting Information for further details).

The remaining fragment-molecule pairs were filtered for several 2D properties, namely synthetic accessibility,^32^ ring aromaticity, and pan-assay interference compounds (PAINS) sub-structures, ^33^ to remove unwanted examples. Full details of the property filters can be found in the Supporting Information.

By filtering the training set for specific 2D properties, we are also able to assess whether the model is able to learn to generate linkers with certain properties implicitly from the data alone. Since these properties are not input explicitly into the model, these could easily be tailored to a specific project or other requirements.

To provide structural information, we generated 3D conformers for the ZINC set using RD-Kit,^34^ adopting the filtering and sampling procedure proposed by Ebejer et al. ^35^. We took the lowest energy conformation as the reference 3D structure for each molecule.

These preprocessing and filtering steps resulted in a data set of 418 797 example fragment elaborations, with linkers of between three and twelve atoms. We selected 800 fragment-molecule pairs at random for model validation (400) and testing (400), and used the remainder to train our model, ensuring no overlap between the molecules in the training and held-out sets.

#### CASF

A major limitation of the ZINC data set is the use of generated comformers, as opposed to experimentally verified active ones. To address this, we used the CASF-2016 data set,^36^ which consists of 285 protein-ligand complexes with high-quality crystal structures from a diverse set of proteins, as an independent test set. We followed the same preprocessing procedure as for the ZINC data set (except for conformer generation), resulting in a set of 309 examples.

We performed large-scale evaluations of our method on both the held-out ZINC test data and the CASF data set. For each example, we generated 250 molecules from each pair of unlinked fragments, assuming the linker length was equal to the linker length of the original molecule.

### Assessment metrics

We assessed the generated molecules with a range of 2D and 3D metrics. As is standard in the assessment of models for molecule generation,^37^ we first checked the generated molecules for validity, uniqueness, and novelty. We then determined if the generated linkers were consistent with the 2D property filters used to produce the training set. In addition, we recorded in how many cases the original molecule used to produce the fragments was recovered by the generation process.

Molecules which passed the 2D property filters were assessed on the basis of their 3D shape. Conformers of the generated molecules and the original molecule were compared using two distinct methods: (i) a shape and colour similarity score (SC_RDKit_), and (ii) root-mean-square deviation (RMSD).

The shape and colour similarity score (SC_RDKit_) uses two RDKit functions, based on the methods described in Putta et al. ^38^ and Landrum et al. ^39^. The colour similarity function scores two 3D conformers against each other based on the overlap of their pharma-cophoric features, while the shape similarity measure is a simple volumetric comparison between the two conformers. Each produces a score between 0 (no match) and 1 (perfect match), which are averaged to produce a final score between 0 and 1. Scores above 0.7 indicate a good match, while scores above 0.9 suggest an almost perfect match. An illustration of several conformers and their similarity scores can be seen in Figure 3.

**Figure 3:**
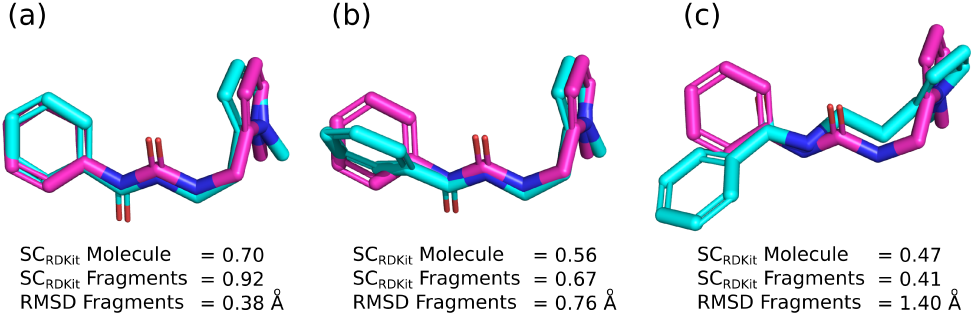
Examples of the 3D metrics used to assess the similarity of conformers. The reference conformer is shown in magenta, while conformers of the generated molecule are shown in cyan. (a) represents very strong alignment by both fragment-based metrics, but lower similarity by SC_RDKit_ Molecule due to the different linker. (b) shows modest similarity by all three metrics, while (c) represents poor similarity by all three measures.

SC_RDKit_ can either be calculated by comparing only the atoms of the starting fragments (SC_RDKit_ Fragments), or by comparing the entire generated molecule to the original molecule (SC_RDKit_ Molecule). The first measure assesses how closely the conformations of the fragments match, whereas the second also incorporates whether or not the generated linker matches the original (Figure 3). Our method is trained to output a diverse range of linkers and not to map exactly to a previously observed linker. However, in the case of scaffold hopping, this metric is important as typically the new linker should match the shape and pharmacophoric features of the original core. ^23^

RMSD between the coordinates of atoms in the starting fragments in the original and generated molecule can be calculated to give a different measure of 3D similarity (RMSD Fragments). A perfect match has an RMSD of 0Å, with a higher figure indicating greater deviation. An RMSD of below 0.5Å suggests an almost perfect match, while an RMSD above 1.0Å corresponds to a poor match given the alignment procedure and number of heavy atoms. An illustration of several conformations and their RMSDs can be seen in Figure 3. Due to the need to match specific atoms, RMSD can only be (reliably) calculated between the atoms of the fragments that are linked, and not the entire molecule.

For each proposed molecule, we generated 3D conformers using RDKit, ^34^ adopting the filtering and sampling procedure proposed by Ebejer et al. ^35^, and scored all conformers. The score for each similarity measure was the best score among all generated conformers for a particular molecule.

## Results and Discussion

We demonstrate DeLinker, a deep generative method that designs a molecule incorporating two starting substructures using 3D structural information. We first checked the impact of the structural information and then assessed our generative method in three experiments: (i) large-scale validation on ZINC (generated conformers), (ii) large-scale validation on CASF (experimentally determined active conformations), and (iii) three case studies covering fragment linking,^40^ scaffold hopping,^41^ and PROTAC design. ^42^

### Importance of structural information

To assess the importance of including structural information, we empirically examined its impact on the generation process (Table 1). We considered three almost identical fragment-molecules pairs based on ZINC7670105 from the held-out ZINC test set (see Methods). In all three cases, the starting substructures remained constant, but the substitution pattern of the benzene linker differed. This resulted in the distance and angle between the fragments changing, but no other differences between the input data to our model. We generated 1000 linkers with a maximum of six atoms for each set of structural information and assessed the substitution pattern of generated molecules that contained a six-membered ring as the linker (Table 1).

**Table 1:**
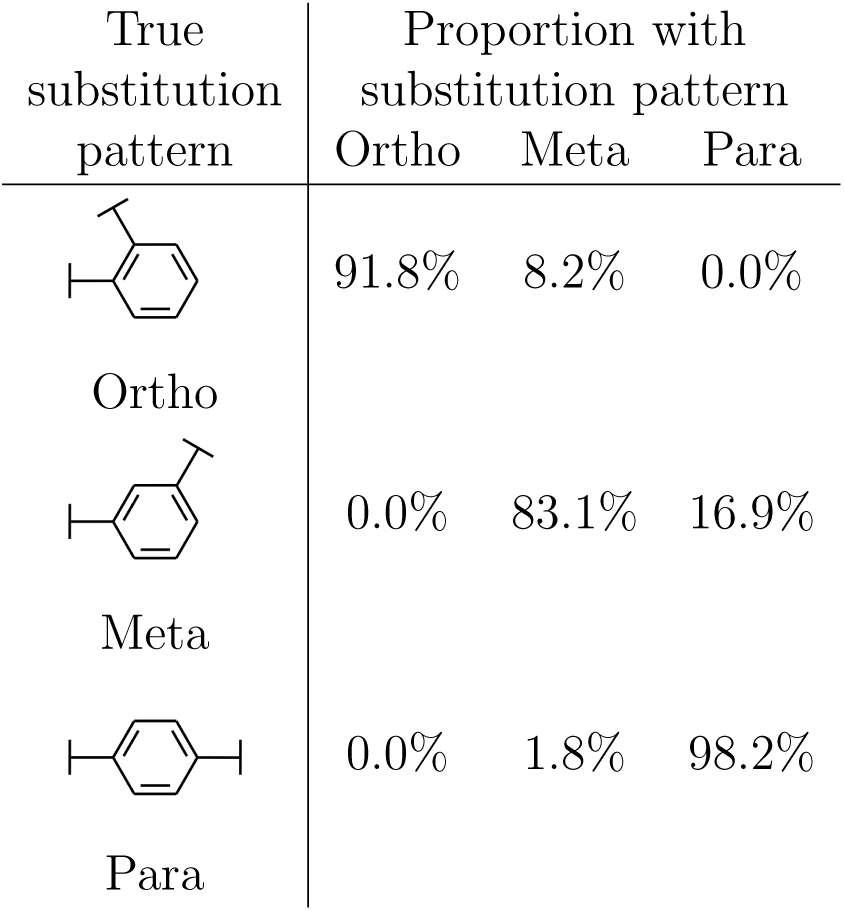
Impact of structural information on generated ring substitution patterns. The generated compounds closely followed the true substitution pattern, with only the structural information provided differing between examples. Importance of structural information provided differing between examples.

DeLinker generated a high number of six-membered rings in all three cases (33%-54%). The most rings were generated with the para-structural information. This is consistent with chemical knowledge since there are fewer possibilities given those structural constraints. The generated molecules closely followed the substitution pattern of the molecule used to calculate the structural information, with between 83% and 98% of the rings produced following the same pattern (Table 1). The effect of the structural information on the performance of DeLinker in a large-scale evaluation is discussed below and can be found in Tables S2 and S3.

### Validation on ZINC

We next evaluated our method on the held-out test set from the ZINC data set, consisting of 400 pairs of fragments. We compared DeLinker to a method based on database lookup (“Database”). The Database samples linkers from the same set of data used to train our method, joining the fragments in one of the two possible orientations at random. This setup ensures that both methods are constructed using the same data, and allows a direct comparison to be made between database lookup and our deep learning-based generative approach.

We generated 250 linkers for each pair of fragments, resulting in 100 000 generated molecules to be assessed for both DeLinker and the Database (see Methods for details). For the evaluation on ZINC, the number of atoms in the linker was set equal to the linker length of the original molecule. This is an easier test for both methods than if the linker length was assumed to be unknown, but allows us to assess whether the two methods presented are able to generate molecules that possess desired 2D chemical properties and high 3D structural similarity.

DeLinker substantially outperformed the Database method by all 3D similarity measures (Table 3), generating a high proportion of valid molecules that passed the 2D chemical property filters (Table 2). Further metrics and an ablation study showing the effects of including different structural information can be found in Tables S2 and S3. Without any structural information, the deep generative model performed similarly to the Database method by the 3D similarity measures (Table S3). Including only the distance between the fragments substantially improved performance, with further benefit from including the angle between fragments.

**Table 2:** 2D performance metrics for molecules generated by DeLinker, our deep generative model, compared to a Database baseline on the held-out ZINC test set and the independent CASF data set.

| Metric | ZINC |  | CASF |  | CASF > 5 atoms |  |
| --- | --- | --- | --- | --- | --- | --- |
|  | Database | DeLinker | Database | DeLinker | Database | DeLinker |
| Valid | 100.0% | 98.4% | 99.0% | 95.5% | 98.4% | 94.7% |
| Unique | 38.8% | 44.2% | 43.0% | 51.9% | 58.3% | 72.9% |
| Novel | 0.0% | 39.5% | 0.0% | 51.0% | 0.0% | 68.7% |
| Recovered | 78.0% | 79.0% | 42.8% | 53.7% | 14.9% | 29.8% |
| Pass 2D filters | 97.0% | 89.8% | 95.0% | 81.4% | 93.6% | 71.7% |

**Table 3:** 3D performance metrics for molecules generated by DeLinker, our deep generative model, compared to a Database baseline on the held-out ZINC test set and the independent CASF data set. See Methods - Assessment metrics for a description of the metrics.

| Metric | ZINC |  | CASF |  | CASF > 5 atoms |  |
| --- | --- | --- | --- | --- | --- | --- |
|  | Database | DeLinker | Database | DeLinker | Database | DeLinker |
| SC<sub>RDKit</sub> Molecule |  |  |  |  |  |  |
| >0.7 | 33.5% | 47.1% | 14.9% | 22.3% | 7.8% | 16.3% |
| >0.8 | 8.5% | 14.2% | 3.3% | 5.2% | 1.1% | 3.6% |
| >0.9 | 1.3% | 1.8% | 0.5% | 0.8% | 0.3% | 0.8% |
| SC<sub>RDKit</sub> Fragments |  |  |  |  |  |  |
| >0.7 | 60.2% | 71.3% | 28.4% | 39.1% | 24.2% | 38.7% |
| >0.8 | 24.7% | 35.8% | 8.7% | 12.7% | 6.1% | 12.3% |
| >0.9 | 4.5% | 8.2% | 1.4% | 2.3% | 0.5% | 1.6% |
| RMSD Fragments |  |  |  |  |  |  |
| <1.00Å | 46.9% | 58.6% | 19.4% | 28.1% | 14.6% | 26.6% |
| <0.75Å | 20.5% | 30.0% | 7.1% | 10.2% | 4.3% | 9.3% |
| <0.50Å | 5.7% | 9.3% | 2.0% | 3.1% | 0.9% | 2.4% |

A molecule is deemed “valid” if it contains both starting fragments (i.e. the fragments have been linked) and its SMILES representation can be parsed by RDKit^34^ (i.e. satisfies atomic valency rules). The small proportion of invalid molecules produced by DeLinker (Table 2) were all due to the fragments remaining unlinked, rather than failing atomic valency. This is a design choice by the deep learning system, and is beneficial in reducing the number of unsuitable linkers suggested.

A fundamental benefit of our deep generative method over any database is evident in the proportion of novel linkers. The Database method is unable to suggest linkers not in the database, and thus 0% of the proposals were novel. In contrast, DeLinker proposed a linker not in the training set in around 40% of suggestions, despite the training set of linkers containing over 5 000 unique linkers. Examples of novel linkers proposed by DeLinker are shown in Figure S1.

Both methods recovered over 75% of the original molecules (Table 2), demonstrating that they are able to sample from the distribution of linkers effectively. However, this is in part due to the chemical redundancy of molecules in ZINC. Indeed, all of the original linkers in the held-out ZINC test set were present in the training set.

DeLinker was able to learn the 2D filters implicitly, although it produced slightly fewer molecules passing these filters than the Database method (Table 2). Both methods had high success rates of 95% or above for all of the individual filters (Table S2).

For all of the 3D measures at all thresholds assessed, DeLinker produced a substantially higher proportion of linkers with the required 3D similarity than the Database (Table 3). In particular, at the highest levels of similarity, DeLinker generated over 80% more molecules scoring > 0.9 for SC_RDKit_ Fragments and over 60% more molecules with an RMSD < 0.5Å.

Performance of both methods is impacted by the length of the generated linkers, and in particular the number of short (three/four atoms) linkers in the test set (Table S1), where there are a limited number of possibilities. The degree of outperformance of DeLinker over the Database increased substantially when only considering linkers with at least five atoms (Table S4). In this setting, DeLinker generated around 190% more molecules scoring > 0.9 for SC_RDKit_ Fragments and 130% more molecules with RMSD Fragments < 0.5Å or SC_RDKit_ Molecule > 0.9.

### Validation on CASF

We saw similar performance when we evaluated the methods on the CASF data set (Tables 2 and 3). Both methods found producing 3D similar molecules more challenging than the held-out ZINC set. However, our method was still frequently able to generate compounds with high similarity to the original molecule (Table 3).

In particular, DeLinker generated around 60% more molecules than the Database at the highest 3D similarity threshold (> 0.9 SC_RDKit_ Fragments and SC_RDKit_ Molecule, < 0.5Å RMSD Fragments). When restricting the evaluation to linkers with at least five atoms, the degree of outperformance substantially increased, with DeLinker producing 200% more molecules that scored > 0.9 by SC_RDKit_ Fragments than the Database (Table 3).

DeLinker recovered 54% of the original linkers, compared to only 43% for the Database method, while around 50% of molecules generated by DeLinker were novel. The proportion recovered was lower than in the evaluation on ZINC, however, this set is more challenging with an average length of the true linker 5.9 atoms, compared to 4.9 for the held-out ZINC test set and 4.7 for the ZINC training set. In addition, only around 70% of the true linkers were present in the training set, providing an upper bound for the Database method. Similarly to the ZINC set, DeLinker substantially outperformed the Database method for longer linkers; DeLinker recovered around 30% of molecules with a linker of at least five atoms, twice as many as the Database method which only recovered 15% (Table 2).

As previously noted, a fundamental limitation of a database method is an inability to generate linkers that are not present in the database. Despite being trained on the same database of linkers, DeLinker has learnt to extrapolate from this set to novel linkers. The following is an example of when this is crucial for successful compound design.

Dequalinium is a nanomolar binder (Ki: 70 nM) of Chitinase A (PDB ID: 3ARP, Figure 4b).^43^ One possible fragmentation of the dequalinium-chitinase complex is shown in Figure 4a. To recover dequalinium from these fragments requires joining them with a decane linker, which is not present in the training set of linkers and thus the Database is unable to recover the original molecule. We generated 250 molecules with DeLinker, which included several highly similar novel linkers. The five most similar by SC_RDKit_ Fragments are shown in Figure 4c. While DeLinker did not recover the decane linker within 250 generated compounds, simple chain linkers that closely resemble the true decane linker are prevalent. We compared this to an exhaustive search of linkers in the Database of the same length as the true decane linker (790 unique molecules). None of the Database generated molecules are highly similar to dequalinium (Figure 4d), with only one molecule with SC_RDKit_ Fragments > 0.7. In contrast, DeLinker generated 34 unique molecules with SC_RDKit_ Fragments > 0.7. This illustrates the importance of *de novo* design and the limitations of any database-based solution.

**Figure 4:**
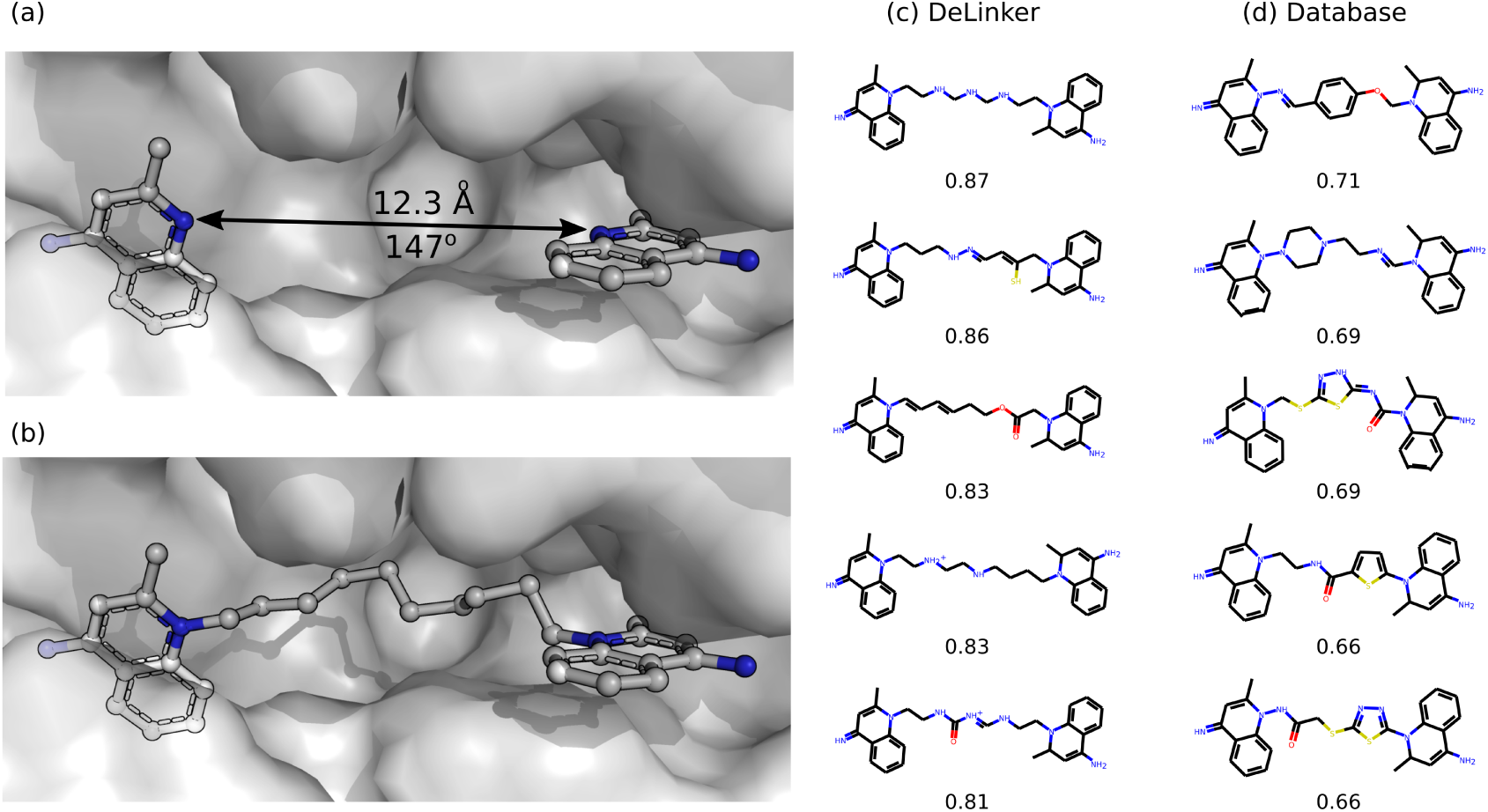
Comparison of DeLinker with an exhaustive Database search. A fragmention of dequalinium (PDB ID: 3ARP, (b)) is shown in (a). The most 3D similar molecules by SC_RDKit_ Fragments proposed by DeLinker and the Database method are shown in (c) and (d), respectively, together with the 3D similarity score. (c) DeLinker was able to produce several very similar molecules, despite limited sampling (250 samples). (d) An exhaustive search of the database was not able to recover the original molecule or produce any highly similar molecules.

Finally, we showed the applicability of our method in three diverse examples from the literature, covering fragment linking, ^40^ scaffold hopping,^41^ and PROTAC design.^42^ Due to the availability of independent experimental structural data for both the initial and optimised complexes, this represents the most realistic evaluation, albeit with a limited sample size.

### Fragment linking case study

Trapero et al. ^40^ considered both growing and linking strategies to create potent inhibitors of inosine 5-monophosphate dehydrogenase (IM-PDH, UniProt: G7CNL4), a tuberculosis drug target. Linking proved most successful, with the authors identifying several promising compounds, the most potent with more than 1000-fold improvement in affinity over the initial fragment hits. Three direct elaborations of the initial fragments were reported (compounds 29-31 in Table 4 of Trapero et al. ^40^), with structures of both the initial fragments (PDB ID: 5OU2) and most potent linked compound (PDB ID: 5OU3) available (Figure 5a).

**Figure 5:**
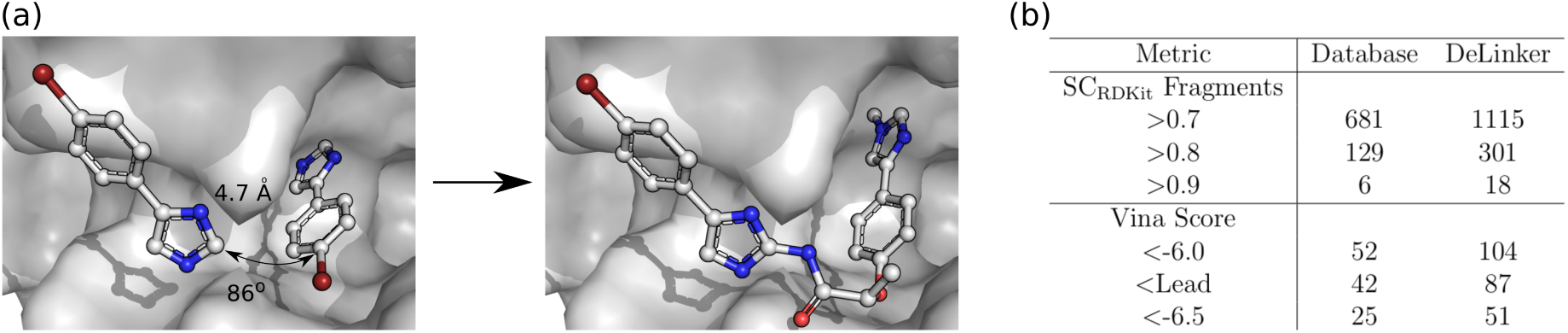
Fragment linking case study. (a) Left: The inital fragment hits (PDB 5OU2). Right: The most potent experimentally verified linked molecule of Trapero et al. ^40^ (PDB ID: 5OU3). DeLinker recovered this and the two other experimentally verified active molecules. (b) 3D similarity metrics and AutoDock Vina Minized Affinities. Unique ligands with SC_RDKit_ Fragments > 0.8 were docked with AutoDock Vina using a local minimization. DeLinker produced more than twice as many molecules than the Database method with better Vina scores than the most potent reported binder (Lead).

In previous experiments, we chose the linker length based on the number of atoms in the linker of the original molecule. To reflect prospective use more accurately, we assumed the linker length was unknown and generated 1000 linkers for each length between three and eleven atoms, inclusively. We assessed the generated linkers using the same criteria as before.

DeLinker recovered all three experimentally validated compounds, while the Database method recovered two, although all three linkers were present in the training set. In addition, DeLinker identified more than twice as many unique compounds as the Database with high 3D similarity (> 0.8 SC_RDKit_ Fragments) to the initial fragments (301 vs. 129, Figure 5b and Table S5). The compounds meeting the above 3D similarity threshold were docked with AutoDock Vina ^44,45^ via a local minimisation after alignment with the starting fragments. This allows us to understand whether the proposed molecules are complementary to the active site and are able to maintain the binding mode of the original fragments. Around 30% of both DeLinker and Database molecules were scored better the most potent experimentally validated compound. As a result, DeLinker suggested more than twice as many unique compounds as the Database with better docking scores than the active compound (87 vs. 42, Figure 5b).

### Scaffold hopping case study

Kamenecka et al. ^41^ designed JNK3-selective (UniProt: P53779) inhibitors that had > 1000-fold selectivity over p38 (UniProt: Q16539), another closely related mitogen-activated protein kinase family member. Starting with an indazole class of compounds, they were not able to establish significant selectivity for JNK3 over p38. However, changing scaffolds led to an aminopyrazole linker that afforded compounds with > 2800-fold selectivity. The two inhibitors displayed nearly identical binding mode (RMSD 0.33 Å, Figure 6a) and affinity for JNK3 (indazole: IC50 12 nM, aminopyrazole: IC50 25 nM), but significantly different binding affinity to p38 (indazole: IC50 3.2 nM, aminopyrazole IC50 3.6 *μ*M).

**Figure 6:**
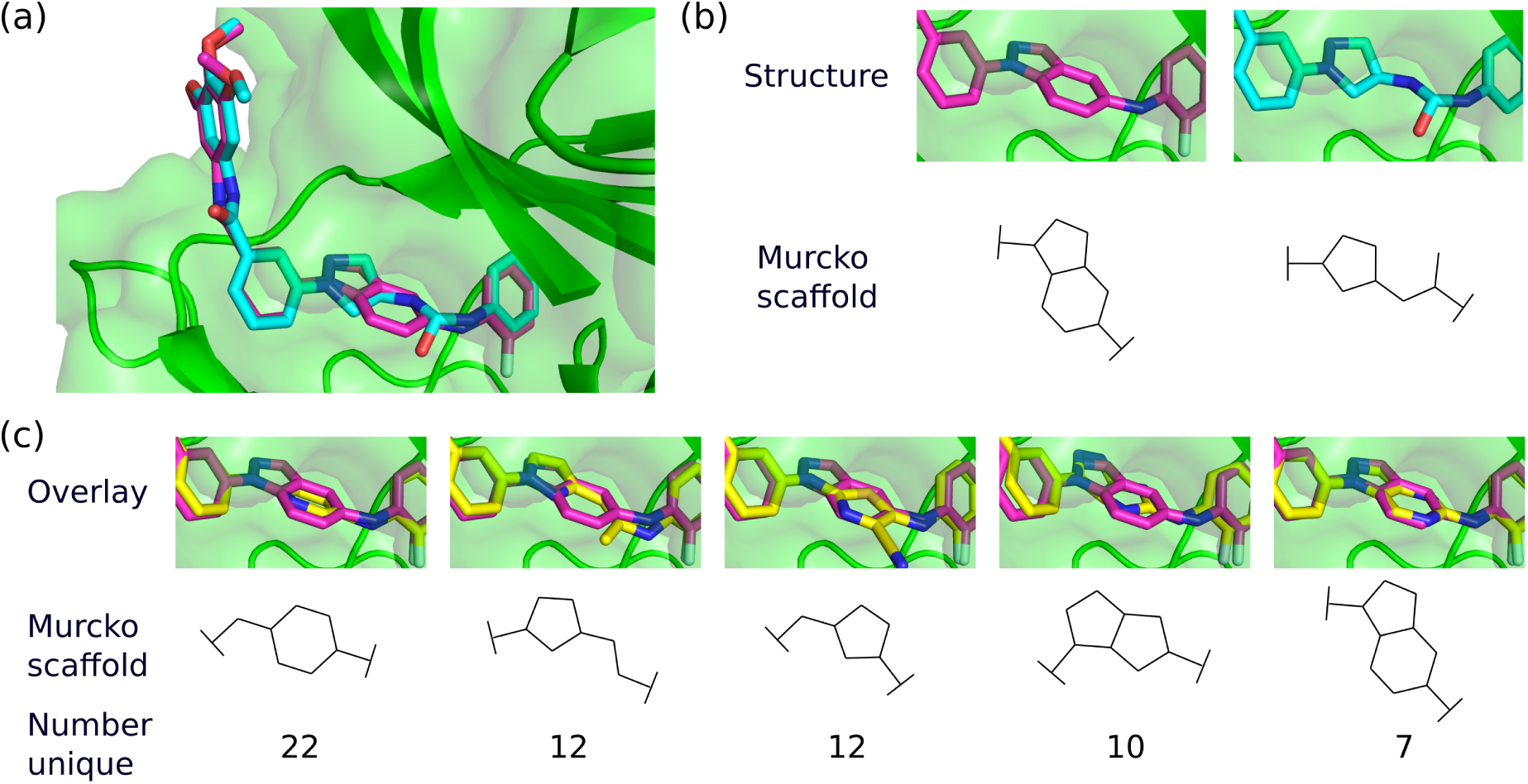
Scaffold hopping case study. (a) Overlay of the indazole (PDB ID 3FI3, magenta carbons) and aminopyrazole (PDB ID 3FI2, cyan carbons) structures, with JNK3 shown in green. DeLinker recovered both active molecules, despite neither linker being in the training set. (b) Structures of the indazole (left) and aminopyrazole (right) linkers, and their Murcko scaffolds. (c) Overlay of the indazole compound (PDB ID 3FI3, magenta carbons) and example linkers (yellow carbons) from several highly 3D similar scaffolds.

Starting with the indazole-based inhibitor (PDB ID: 3FI3), we explored the ability of our method to change molecular scaffold, in particular towards the aminopyrazole-based inhibitor (PDB ID: 3FI2). We generated 5 000 linkers with both eight and nine atoms, and assessed the generated linkers using the same criteria as before. In particular, we focussed on the diversity of molecular scaffolds proposed by DeLinker that satisfied the 3D structural information and could adopt a highly similar conformation to the original indazole-based inhibitor.

Of the 10 000 compounds generated by DeLinker, there were 2 688 unique compounds that satisfied the 2D chemical filters (Table S6). 699 of these had a SC_RDKit_ Fragments score above 0.75, of which 627 were not in the training set (89.7% novel). These compounds covered 182 unique generic Murko scaffolds. ^46^ Five of the most common are shown in Figure 6b, together with an example linker and the number of unique linkers generated with the same generic Murko scaffold that also met the 3D similarity threshold. The examples from all five scaffolds show almost perfect overlap with the indazole linker, while maintaining the conformation of the remainder of the molecule. In addition, DeLinker recovered both the indazole- and aminopyrazole-based linkers, despite neither being present in the training set.

### PROTAC case study

Farnaby et al. ^42^ developed PROTAC degraders of the BAF AT-Pase subunits SMARCA2 (UniProt: P51531) and SMARCA4 (UniProt: P51532) using a bromodomain ligand and recruitment of the E3 ubiquitin ligase VHL (UniProt: P40337). They first designed a PROTAC by combining known binders of SMARCA2/4 and E3 ubiquitin ligase VHL using polyethylene glycol-based linkers (PDB ID: 6HAY, Figure 7a). The linker was then optimised to improve interactions with the lipophilic face created in part by Y98 of the VHL protein. In particular, they designed the linker to mimic the conformation observed in the ternary complex structure, resulting in improved molecular recognition (PDB ID: 6HAX). This was confirmed with the two crystal structures displaying near identical ternary complexes (Figure 7b).

**Figure 7:**
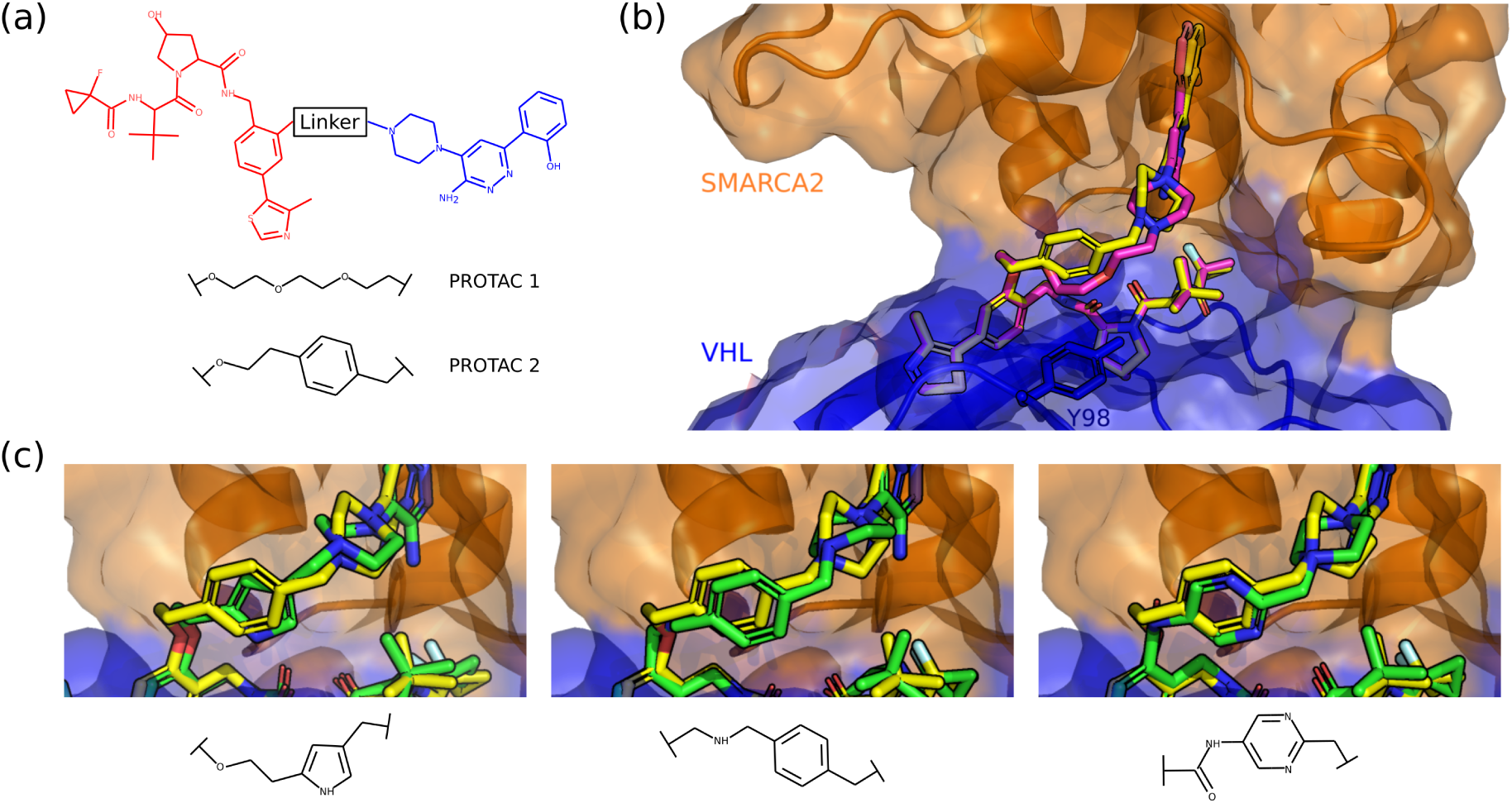
PROTAC design case study. (a) Two-dimensional chemical structures of PROTAC 1 and PROTAC 2. (b) Overlays of ternary crystal structures of PROTAC 1 (PDB ID 6HAY, magenta carbons) and PROTAC 2 (PDB ID 6HAY, yellow carbons), with SMARCA2 shown in orange, VHL in blue. (c) Overlays of three linkers with different scaffolds produced by DeLinker (green carbons); all three accurately recapitulate the linker geometry observed in PROTAC 2 (yellow carbons). None of these linkers were present in the training set.

We investigated the ability of our model to design alternative linkers to the known polyethylene glycol-based linker (PDB ID: 6HAY) that could maintain the same conformation observed in the ternary complex. We generated 5 000 linkers with a maximum of either nine or ten atoms. There were almost 3 000 unique linkers that passed the 2D chemical filters (Table S7).

Due to the size and complexity of the PROTAC, we generated conformers constraining the two starting substructures (Figure 7a) to adopt poses close to their known binding conformation, removing any high energy poses. DeLinker produced 236 unique compounds with SC_RDKit_ Fragments > 0.85, of which three novel linkers that accurately recapitulate the linker geometry observed in PROTAC 2 are shown in Figure 7c. In all three cases, the aromatic systems perfectly align with that of PROTAC 2, and are likely to fulfil the goal of improving interactions with the lipophilic face compared to PROTAC 1. In particular, the pyrrole-based linker (Figure 7c, left) appears to be making a NH-*π* interaction with the Y98 residue, improving the CH-*π* interaction being made by the benzene in PROTAC 2.

## Conclusion

We have developed a graph-based deep generative method for fragment linking or scaffold hopping that is protein context dependent, utilising the relative distance and orientation between the starting substructures in the design process. Unlike previous attempts at computational fragment linking or scaffold hopping, our method does not rely on a database of fragments from which to select a linker but instead designs one given the fragments provided and 3D information.

Through two large scale assessments, we have demonstrated that our generative method is able to learn to produce a distribution of linkers that matches the constraints present in the training set, while being able to generalise to novel linkers that satisfy both 2D and 3D constraints. In addition, the generated molecules consistently have high 3D similarity to both the initial fragments and the original molecules, outperforming a database baseline by 60% in the evaluation on CASF, increasing to 200% when restricting the evaluation to linkers with at least five atoms.

Finally, through three case studies, we have shown that our method can be applied to fragment linking, scaffold hopping, and PROTAC design. In the fragment linking example, our method reproduced all of the reported potent molecules using only the crystal data of the initial fragment hits. In addition, in docking-based evaluation, many of the generated molecules were scored more highly than the original hits, while maintaining similar binding modes. In the scaffold hopping case study, our method reproduced both the starting and final molecule, while suggesting many other scaffolds with high 3D similarity to the initial crystal data. Finally, in the PROTAC design case study, our method suggested a range of novel linkers that met the design goal of maintaining the linker geometry of PROTAC 1, while improving interactions with the lipophilic face created in part by residue Y98 of the VHL protein.

As far as we are aware, this is the first molecular generative model to incorporate 3D structural information directly in the design process. Currently the only 3D information utilised by the model is the distance between the fragments or starting substructures and their relative orientations. This provides explicit constraints for a given compound, but only implicit information about the shape of the binding site. Despite this minimal parametrisation, there is a substantial impact on the generated molecules. Extending our method to use additional structural information that incorporates further constraints from the protein is a direction for future research and promises substantial benefits for structure-based molecular generative methods.

Code is available at https://github.com/oxpig/DeLinker.

## Acknowledgement

The authors thank Veerabahu Shanmugasundaram for helpful discussions on PROTACs. F.I. is supported by the EPSRC (Reference: EP/N509711/1).

