## Supporting Information for "Deep Generative Models for 3D Compound Design"

### Supporting Information
### Deep Generative Models for 3D Compound Design

Fergus Imrie,[†] Anthony R. Bradley,[‡] Mihaela van der Schaar,[¶,§] and Charlotte M. Deane[*,†]

†*Oxford Protein Informatics Group, Department of Statistics, University of Oxford, Oxford OX1 3LB, UK*

‡*Exscientia Ltd, 36 St. Giles', Oxford, OX1 3LD, UK*

¶*University of Cambridge, Cambridge CB2 1PZ, UK*

§*Alan Turing Institute, London NW1 2DB, UK*

### DeLinker Implementation Details

#### Atom types.

There are 14 permitted atom types: carbon, nitrogen ($N^-$, N, $N^+$), oxygen ($O^-$, O, $O^+$), fluorine, chlorine, bromine, iodine, and sulphur (maximum valence 2, 4, or 6).

#### Network architecture.

Both the encoder and decoder are standard gated graph neural networks (GGNN),[1] which propagate messages for 7 steps, and have residual connections between odd numbered time steps.

We implemented the function $f$, which maps the hidden state of a node to its atom type, as a linear classifier with attention from the node's hidden vector to one of the node types. The attention mechanism is similar to Bahdanau et al.[2] and allows the label for a given node to depend on the hidden states of the other expansion nodes.

We trained the model with a learning rate of 0.001 for 10 epochs using the Adam optimiser.

#### Hyperparameter search.

We performed a limited hyperparameter search of the following parameters (final parameters in bold):

- Learning rate: 0.01, **0.001**, 0.0001
- Batch size: 8, **16**,
- Hidden state dimension: 16, **32**, 50, 100
- Encoding dimension: 2, **4**, 8
- $\lambda_{KL}$: 0.1, **0.3**, 0.6

The model was fairly robust to the choice of hyperparameters. Performance was measured via the validation reconstruction loss and not generative performance.

### Data curation.

Fragment-molecule pairs for the ZINC[3] and CASF[4] sets were constructed as follows. First all possible fragmentations of each molecule were produced by enumerating all double acyclic single bond cuts.[5] These we then filtered to remove trivial and unrealistic situations using the following constraints: (i) minimum linker length: 3 atoms, (ii) minimum fragment size: 5 atoms, (iii) linker fewer heavy atoms than either fragment, (iv) minimum path length between fragments: 2 atoms.

The remaining fragment-molecule pairs were filtered for several 2D properties: (i) the synthetic accessibility (SA) score[6] of the molecule must be lower than the fragments with exit vectors represented by dummy atoms, (ii) the molecule must pass pan-assay interference (PAINS)[7] filters, and (iii) rings must either be saturated aliphatic or aromatic (according to RDKit[8] valency rules). PAINS filters were implemented by SMARTS substructure searching with RDKit, using the RDKit version of the Saubern et al.[9] translation of the original PAINS.[7] as specified at `https://github.com/rdkit/rdkit/blob/master/Data/Pains/wehi_pains.csv` (Accessed: 02/06/2019). Any molecules containing atom types outside of the permitted atom types were excluded.

### Training set composition

Table S1: Distribution of number of atoms contained in the original linkers in the datasets utilised. The average linker length in the CASF set (5.9) is around one atom longer than the ZINC training set (4.7), validation (4.7) and test set (4.9).

| Linker | ZINC | | | CASF |
|---|---|---|---|---|
| Length | Train | Valid | Test | Test |
| 3 | 28.7% | 30.7% | 26.2% | 25.2% |
| 4 | 21.6% | 22.7% | 17.7% | 13.9% |
| 5 | 19.5% | 15.7% | 22.0% | 10.7% |
| 6 | 17.8% | 16.5% | 21.0% | 13.3% |
| 7 | 7.7% | 10.3% | 7.3% | 12.0% |
| 8 | 3.1% | 2.5% | 3.5% | 9.1% |
| 9 | 1.3% | 1.3% | 2.0% | 4.8% |
| 10 | 0.3% | 0.3% | 0.3% | 3.2% |
| 11 | 0.0% | - | - | 5.5% |
| $\geq$12 | 0.0% | - | - | 2.3% |

### Additional results

Table S2: Ablation study for DeLinker, our deep generative method on the ZINC data set. We show the effect on the 2D metrics of removing all of the structural information ("No info") and including only the distance information ("Distance") compared to our full protocol ("DeLinker") and the database baseline ("Database"). See Data curation for a description of the 2D property filters.

| Metric | Database | No Info | Distance | DeLinker |
|---|---|---|---|---|
| Valid | 100.0% | 97.0% | 98.6% | 98.4% |
| Unique | 38.8% | 51.2% | 47.3% | 44.2% |
| Novel | 0.0% | 36.2% | 37.6% | 39.5% |
| Recovered | 78.0% | 74.5% | 78.3% | 79.0% |
| Pass 2D filters | 97.0% | 89.9% | 90.2% | 89.8% |
| Pass SA filter | 97.8% | 95.1% | 95.5% | 95.3% |
| Pass ring filter | 100.0% | 95.2% | 94.5% | 94.8% |
| Pass PAINS filter | 99.2% | 97.8% | 98.4% | 97.9% |

Table S3: Ablation study for DeLinker, our deep generative method, on the ZINC data set. We show the effect on the 3D metrics of removing all of the structural information ("No info") and including only the distance information ("Distance") compared to our full protocol ("DeLinker") that includes both distance and angle information, and the database baseline ("Database"). See Methods - Assessment metrics for a description of the metrics.

| Metric | Baseline | No Info | Distance | DeLinker |
|---|---|---|---|---|
| $SC_{RDKit}$ Molecule | | | | |
| >0.7 | 35.5% | 37.6% | 43.2% | 47.1% |
| >0.8 | 8.5% | 9.2% | 11.8% | 14.2% |
| >0.9 | 1.3% | 1.1% | 1.5% | 1.8% |
| $SC_{RDKit}$ Fragments | | | | |
| >0.7 | 60.2% | 64.4% | 69.1% | 71.3% |
| >0.8 | 24.7% | 27.7% | 33.4% | 35.8% |
| >0.9 | 4.5% | 5.0% | 7.0% | 8.2% |
| RMSD Fragments | | | | |
| <1.00 | 46.9% | 50.9% | 56.6% | 58.6% |
| <0.75 | 20.5% | 22.4% | 27.8% | 30.0% |
| <0.50 | 5.7% | 5.6% | 7.9% | 9.3% |

Table S4: 2D and 3D metrics for molecules generated by DeLinker, our *de novo* deep generative model, compared to a Database baseline on the held-out ZINC test set. See Data curation for a description of the 2D property filters and Methods - Assessment metrics for a description of the 3D metrics.

| | ZINC | | ZINC > 5 atoms | |
|---|---|---|---|---|
| Metric | Database | DeLinker | Database | DeLinker |
| Valid | 100.0% | 98.4% | 99.0% | 95.5% |
| Unique | 38.8% | 44.2% | 43.0% | 51.9% |
| Novel | 0.0% | 39.5% | 0.0% | 51.0% |
| Recovered | 78.0% | 79.0% | 42.8% | 53.7% |
| Pass 2D filters | 97.0% | 89.8% | 95.0% | 81.4% |
| $SC_{RDKit}$ Molecule | | | | |
| >0.7 | 33.5% | 47.1% | 21.3% | 37.1% |
| >0.8 | 8.5% | 14.2% | 3.5% | 9.4% |
| >0.9 | 1.3% | 1.8% | 0.4% | 1.0% |
| $SC_{RDKit}$ Fragments | | | | |
| >0.7 | 60.2% | 71.3% | 51.5% | 66.7% |
| >0.8 | 24.7% | 35.8% | 16.8% | 30.3% |
| >0.9 | 4.5% | 8.2% | 2.1% | 6.0% |
| RMSD Fragments | | | | |
| <1.00Å | 46.9% | 58.6% | 39.1% | 55.1% |
| <0.75Å | 20.5% | 30.0% | 14.2% | 26.9% |
| <0.50Å | 5.7% | 9.3% | 3.0% | 6.9% |

Table S5: Fragment linking case study. 2D and 3D Metrics for DeLinker and the Database baseline.

| Metric | Database | DeLinker |
|---|---|---|
| Valid | 100.0% | 98.7% |
| Unique | 30.7% | 56.4% |
| Novel | 0.0% | 58.5% |
| Recovered | 66.7% | 100.0% |
| Pass 2D filters | 97.8% | 74.0% |
| $SC_{RDKit}$ Fragments | | |
| >0.7 | 681 | 1115 |
| >0.8 | 129 | 301 |
| >0.9 | 6 | 18 |

Table S6: Scaffold hopping case study. 2D and 3D metrics for DeLinker.

| Metric | DeLinker |
|---|---|
| Valid | 99.5% |
| Unique | 63.7% |
| Novel | 88.5% |
| Recovered | 100.0% |
| Pass 2D filters | 51.4% |
| $SC_{RDKit}$ Fragments | |
| >0.70 | 1928 |
| >0.75 | 699 |
| >0.80 | 114 |
| >0.85 | 9 |

Table S7: PROTAC design case study. 2D and 3D metrics for DeLinker.

| Metric | DeLinker |
|---|---|
| Valid | 96.0% |
| Unique | 62.1% |
| Novel | 95.9% |
| Recovered | 0.0% |
| Pass 2D filters | 61.8% |
| $SC_{RDKit}$ Fragments | |
| >0.70 | 2926 |
| >0.75 | 2883 |
| >0.80 | 2584 |
| >0.85 | 236 |

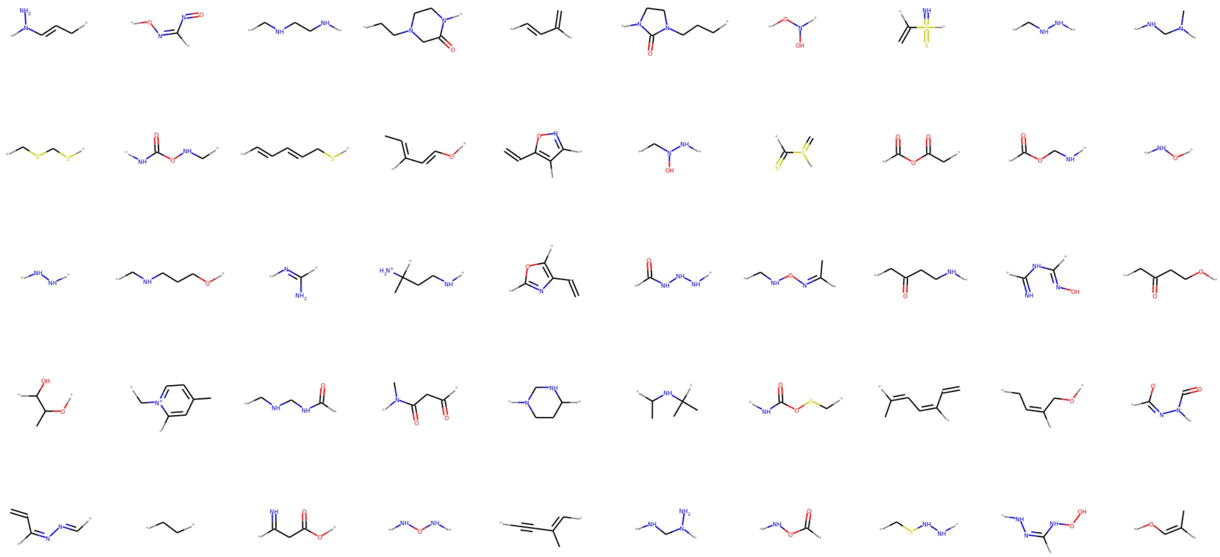

Figure S1: A random sample of 50 novel linkers generated by DeLinker during testing on the held-out ZINC data set.

(6) Ertl, P.; Schuffenhauer, A. Estimation of synthetic accessibility score of drug-like

molecules based on molecular complexity and fragment contributions. *J. Cheminf.* **2009**, *1*, 8.

(7) Baell, J. B.; Holloway, G. A. New Substructure Filters for Removal of Pan Assay Interference Compounds (PAINS) from Screening Libraries and for Their Exclusion in Bioassays. *J. Med. Chem* **2010**, *53*, 2719–2740.

(8) Landrum, G. RDKit: Open-source cheminformatics. `http://www.rdkit.org/`.

(9) Saubern, S.; Guha, R.; Baell, J. B. KNIME Workflow to Assess PAINS Filters in SMARTS Format. Comparison of RDKit and Indigo Cheminformatics Libraries. *Mol. Inf.* **2011**, *30*, 847–850.
